# Inter-laboratory automation of the *in vitro* micronucleus assay using imaging flow cytometry and deep learning

**DOI:** 10.1101/2021.05.05.442619

**Authors:** John W. Wills, Jatin R. Verma, Benjamin J. Rees, Danielle S. G. Harte, Qiellor Haxhiraj, Claire M. Barnes, Rachel Barnes, Matthew A. Rodrigues, Minh Doan, Andrew Filby, Rachel E. Hewitt, Catherine A. Thornton, James G. Cronin, Julia D. Kenny, Ruby Buckley, Anthony M. Lynch, Anne E. Carpenter, Huw D. Summers, George Johnson, Paul Rees

## Abstract

The *in vitro* micronucleus assay is a globally significant method for DNA damage quantification used for regulatory compound safety testing in addition to inter-individual monitoring of environmental, lifestyle and occupational factors. However it relies on time-consuming and user-subjective manual scoring. Here we show that imaging flow cytometry and deep learning image classification represents a capable platform for automated, inter-laboratory operation. Images were captured for the cytokinesis-block micronucleus (CBMN) assay across three laboratories using methyl methanesulphonate (1.25 – 5.0 µg/mL) and/or carbendazim (0.8 – 1.6 µg/mL) exposures to TK6 cells. Human-scored image sets were assembled and used to train and test the classification abilities of the “DeepFlow” neural network in both intra- and inter-laboratory contexts. Harnessing image diversity across laboratories yielded a network able to score unseen data from an entirely new laboratory without any user configuration. Image classification accuracies of 98%, 95%, 82% and 85% were achieved for ‘mononucleates’, ‘binucleates’, ‘mononucleates with MN’ and ‘binucleates with MN’, respectively. Successful classifications of ‘trinucleates’ (90%) and ‘tetranucleates’ (88%) in addition to ‘other or unscorable’ phenotypes (96%) were also achieved. Attempts to classify extremely rare, tri- and tetranucleated cells with micronuclei into their own categories were less successful (≤ 57%). Benchmark dose analyses of human or automatically scored micronucleus frequency data yielded quantitation of the same equipotent dose regardless of scoring method. We conclude that this automated approach offers significant potential to broaden the practical utility of the CBMN method across industry, research and clinical domains. We share our strategy using openly-accessible frameworks.

## INTRODUCTION

Across industry, government and academic research institutions the *in vitro* micronucleus test is one of the most widely used bioassays for the identification and quantification of chromosomal damage (Decordier and Kirsch-Volders 2006; Fenech 2000; Fenech 2020; Kirsch-Volders et al. 2011). Because DNA damage at the chromosome level is recognised as a key event in the initiation of carcinogenesis, the assay has become an essential component of genetic toxicity screening programmes worldwide (Fenech 2000). Harmonised assay protocols and scoring approaches have been detailed by Organisation for Economic Cooperation and Development (OECD)-Test Guideline 487 (OECD 2016). In addition to regulatory compound screening, the assay is also widely used for more specific research and clinical purposes including compound mode-of-action determinations, tumour radiosensitivity prediction and inter-individual monitoring of lifestyle, occupational and environmental factors including radiation biodosimetry assessments (Decordier and Kirsch-Volders 2006; Fenech 2000; Fenech 2020; Kirsch-Volders et al. 2011; Wang et al. 2019).

The micronucleus assay operates through the detection of whole chromosomes or chromosome fragments expressed by cells after nuclear division as satellite ‘micronucleus’ (MN) events. Because complete nuclear division is required to enable expression of these events, the ‘cytokinesis-block’ version of the assay was developed. This method inhibits cell division into daughter entities (cytokinesis) using the microfilament assembly inhibitor cytochalasin-B. This yields cells that have successfully undergone division easily identifiable by their binucleated appearance. In this way, the cytokinesis-block micronucleus (CBMN) assay allows scoring of micronucleus events in cells known to have undergone division during the treatment period. This avoids misleading results otherwise present due to pre-existing damage, sub-optimal cell culture conditions or from the selection of overly cytotoxic compound doses that retard or inhibit cell division and concomitant micronucleus expression (Decordier and Kirsch-Volders 2006; Fenech 2000; Kirsch-Volders et al. 2011). Despite almost global utilisation, CBMN assay scoring still often relies upon manual observation and recording using light microscopy. Whilst manual scoring is the ‘gold standard’, it suffers from user subjectivity and scorer variability in addition to being extremely time and labour-intensive (Rodrigues et al. 2014a; Rodrigues et al. 2014b; Rodrigues et al. 2018). For these reasons, over the last two decades significant efforts have been directed towards automated approaches for both image collection and subsequent scoring. As recently reviewed (Rodrigues et al. 2018), these largely involve slide and laser scanning microscopy systems that automate image collection in conjunction with traditional, threshold-based image classification techniques (Darzynkiewicz et al. 2011; Decordier et al. 2009; Decordier et al. 2011; François et al. 2014; Maertens and White 2015; Rossnerova et al. 2011; Schunck et al. 2004; Seager et al. 2014; Smolewski et al. 2001; Varga et al. 2004; Verhaegen et al. 1994; Willems et al. 2010). Conventional flow cytometry methods have also been developed that aim to identify isolated micronuclei using fluorescence intensity measurements in the absence of image-based validation (Avlasevich et al. 2006; Bryce et al. 2008; Bryce et al. 2010; Bryce et al. 2013; Bryce et al. 2007).

More recently, imaging flow cytometry unites the acquisition approach of flow cytometry with microscopical observation (Allemang et al. 2021; Rodrigues 2018; Rodrigues 2019; Rodrigues et al. 2014a; Rodrigues et al. 2014b; Rodrigues et al. 2016a; Rodrigues et al. 2018; Rodrigues et al. 2016b; Wang et al. 2019; Wilkins et al. 2017). This fluidics-based approach is well suited for processing cell suspension cultures (*e*.*g*., TK6 B-lymphocytes commonly used for the CBMN assay) enabling rapid collection of transmitted light brightfield, darkfield laser scatter and fluorescence images for populations of tens of thousands of single cells. Simple inclusion of a single nuclear fluorescent stain (*e*.*g*., Hoechst 33342, propidium iodide or DRAQ5 *etc*.) allows detection of parent nuclei and micronucleus events (Rodrigues 2018; Rodrigues 2019; Rodrigues et al. 2018; Rodrigues et al. 2016b). Without need of further labels, the brightfield images provide essential context for detecting micronuclei associated with parent cells (Rodrigues et al. 2014a; Verma et al. 2018). The ‘Amnis ImageStream^X^’ series cytometers (Luminex Corporation) further support unassisted data acquisition for multiple samples via a 96-well plate sampling attachment. Images are stored to sample-specific data files enabling archiving should human validation or reevaluation be required (Rodrigues et al. 2018). Traditional image classification approaches deployed within the manufacturer-supplied analysis software have shown utility for CBMN scoring automation (Rodrigues 2018; Rodrigues 2019; Rodrigues et al. 2014a; Rodrigues et al. 2014b; Rodrigues et al. 2016a; Rodrigues et al. 2018; Rodrigues et al. 2016b; Wang et al. 2019; Wilkins et al. 2017). However, in our experience these strategies require significant expertise to set up, in addition to frequent tuning to maintain acceptable performance, even within a single laboratory (Verma et al. 2018). Deviations of around 30% from the results obtained by manual microscopy scoring have also been reported in experiments utilising this approach to study irradiated peripheral blood lymphocytes (Rodrigues et al. 2016b). This outcome was in part attributed to the lack of flexibility of the implemented image analysis algorithms relative to the expertise of human judgement (Rodrigues et al. 2018; Rodrigues et al. 2016b).

Building image classification strategies that *generalise* well enough to permit robust, entirely automated image classifications without need of human intervention or configuration is a difficult task. This is because, even when protocols are harmonised, there will always be variability (*e*.*g*., illumination, focus and fluorescence staining heterogeneity *etc*.) in the input image data. This variation is even more extreme across laboratories due to the inevitable use of different imaging equipment, calibration settings, personnel, cell culture and bioassay regimens. Recently, artificial intelligence approaches have been achieving increasing success in providing generalised automation of image classification tasks (Caicedo et al. 2019; Moen et al. 2019). These approaches can use handcrafted features extracted from images in conjunction with machine learning algorithms, but increasingly, the availability of computational power is enabling the application of deep learning on image pixel data (Blasi et al. 2016; Eulenberg et al. 2017). This approach uses so-called deep convolutional neural networks in a manner inspired by neural connectivity in the brain. A typical image classification workflow involves assigning ‘ground truth’ class annotations to a large set of images before subdividing them into ‘train’ and ‘test’ datasets. The weights connecting the nodes of the neural network are then optimised during a training phase that attempts to match the input images to the annotated classifications. A potential issue due to the flexibility of neural networks as non-linear function approximators is that ‘memorisation’ due to over-fitting of training data can emerge (Zhang et al. 2017). For this reason, final network accuracy is assessed by cross validation against a test set that importantly was entirely ‘unseen’ during the training phase. Subsequently, the trained neural net can be deployed for the classification of new images.

In the context of the CBMN assay, deep learning approaches were recently used on imaging flow cytometry data using the cytometer manufacturer’s ‘Amnis Artificial Intelligence’ software to identify binucleated cells in the 3-D reconstructed skin micronucleus assay. This binucleated cell population was then used as a refined start point from which to expedite manual identification of micronucleus events (Allemang et al. 2021). However, there would be considerable value in openly accessible frameworks for accessibility and for adaptability: the modular nature of modern, open source deep learning interfaces allows new network architectures to be easily switched or specifically tailored as they emerge. This flexibility provides complete ability to build bespoke solutions using the latest tools to pursue maximal accuracy and the accommodation of diverse research objectives.

Here, we used imaging flow cytometry to automate image capture for the CBMN assay across three laboratories using differing local protocols for cell culture, bioassay procedure, DNA staining, cytometer calibration and image collection. Given the inherent variability in the captured images, we investigate the ability of deep learning to enable robust, inter-laboratory scoring automation. To do this, we provide an open framework that utilises the powerful, yet lightweight DeepFlow neural network architecture that has been previously optimised to achieve rapid training and classification of imaging flow cytometry data (Eulenberg et al. 2017).

## MATERIALS & METHODS

### Multi-centre image collection

Image data was collected using three different Amnis ImageStream^X^ imaging flow cytometers (Luminex Corporation, USA) across three locations: Central Biotechnology Services, Cardiff University School of Medicine (hereafter, Cardiff), the Department of Veterinary Medicine’s Imaging Facility, University of Cambridge, UK (Cambridge) and at GlaxoSmithKline Research and Development, Stevenage, UK (GSK).

#### Chemicals

Methyl methanesulphonate (MMS) (#129925) (CAS registry number 66-27-3) and carbendazim (#378674) (CAS no. 10605-21-7) were purchased from Sigma-Aldrich (Merck), UK.

#### Cardiff and Cambridge: Cell culture and cytokinesis-block micronucleus assay

P53 competent, virally transformed human B lymphoblastoid (TK6) cells were purchased from the Health Protection Agency Culture Collections (Wiltshire, UK). The cells were cultured in RPMI 1640 media (#A1049101, ThermoFisher) supplemented with 100 U/mL penicillin and 100 µg/mL streptomycin and containing 10% (v/v) heat-inactivated horse serum (#26050088, ThermoFisher). Cells were seeded at 2 × 10^5^ cells/mL in 25 cm^2^ flasks (ThermoFisher) and incubated at 37 °C for ∼ 1.5 cell cycles (24-30 h) in the presence of MMS (0 / 1.25 / 2.5 / 5.0 µg/mL doses) or carbendazim (0 / 0.8 / 1.0 / 1.6 µg/mL doses) with co-exposed cytochalasin-B (#C6762, Sigma) added to a final concentration of 3 µg/mL as a cytokinesis-block. Following exposure, cells were pelleted by centrifugation (200xg, 10 min) and washed once with 10 mL phosphate buffered saline (PBS). Cells were then pelleted and resuspended in 2 mL 1X BD FACS lysing solution (#349202, BD) for 12 min to achieve fixation and permeabilisation.

#### GSK: Cell culture and cytokinesis-block micronucleus assay

TK6 (IVGT) cells (#13051501) purchased from ECACC, operated by Public Health England (Wiltshire, UK). The cells were cultured in RPMI 1640 media with 2 mM glutamine (#52400-025, ThermoFisher) supplemented with 100 U/mL penicillin and 100 µg/mL streptomycin (#15140-122, ThermoFisher), 1.8 mM sodium pyruvate (#11360-039, ThermoFisher) and containing 10% (v/v) heat-inactivated horse serum (#26050-088, BioSera, Labtech, UK). Cells were seeded at 2 × 10^5^ cells/mL in 25 cm^2^ flasks (ThermoFisher) and incubated at 37 °C for 24 h in the presence of carbendazim (0 / 0.8 / 1.2 / 1.6 µg/mL doses) with co-exposed cytochalasin-B (#C6762, Sigma) added to a final concentration of 6 µg/mL as a cytokinesis-block. Following exposure, cells were pelleted by centrifugation (200xg, 10 min) and washed once with 10 mL PBS (#10010-015, ThermoFisher). Cells were then pelleted and resuspended in 2 mL 1X BD FACS lysing solution (#349202, BD) for 12 min to achieve fixation and permeabilisation.

### Nuclear labelling

Fixed, permeabilised cells were incubated with nuclear stains in PBS at room temperature. Nuclei and micronuclei were stained at the Cardiff and GSK laboratories by 30 min incubation with 0.05 mM DRAQ5 (peak excitation: 647 nm, peak emission: 681 nm) (#564902, BD). Samples at the Cambridge laboratory were stained with a 1:2500 dilution (8 µM) of Hoechst 33342 (peak excitation: 351 nm, peak emission: 461 nm) (#62249, ThermoFisher) for 30 mins. After labelling, cells were pelleted, resuspended and final cell concentrations adjusted through addition of PBS towards an optimal cell concentration for imaging flow cytometry (typically ∼100 µL sample volumes at ∼10^7^ cells/mL).

### Imaging flow cytometry

Brightfield and nuclear fluorescence images (20,000 images / sample) were collected using Amnis ImageStream^X^ (Luminex) flow cytometers using the 40X objective lens via the manufacturer’s INSPIRE software at the Cardiff, Cambridge and GSK laboratories (described above). At Cardiff and GSK, DRAQ5-labelled cells were excited using 488 nm or 642 nm lasers (respectively) with the brightfield collected in channel 1 and DRAQ5 in channel 11. At Cambridge, Hoechst 33342-labelled cells were excited using a 405 nm laser with brightfield collection in channel 4 and nuclear fluorescence collection in channel 1. At all locations, a brightfield area range of 100-900 µm^2^ was used to avoid debris, speed bead and large aggregate image collection. Full details of image acquisition settings including the laser excitation powers the exact cytometer models utilised at each location are provided in **Supp. Table S1**.

### Compensated image file generation using IDEAS

Prior to image extraction, raw image files (.rif) acquired by the INSPIRE software were converted to compensated image files (.cif) using identical settings via batch processing with a template using the IDEAS (version 6.2) software (Luminex). During the process, populations of cell images suitable for scoring were refined by gating out (brightfield area, 200 – 500 µm^2^ versus aspect ratio, 0.75 – 1.0) debris and identifying a single cell population that was also suitably in focus. This was achieved by linescan gradient via the root mean square of the brightfield images ranging from 55 – 80.

### Image data pre-processing: CIF to TIF extraction

Single, in-focus cell populations were exported from the IDEAS software in compensated image file format (.cif). The individual cell images within these files were then extracted to 16-bit grayscale, two-channel (nuclear fluorescence / brightfield) multipage TIF files using a custom script (code and example available for download from the BioStudies database (http://www.ebi.ac.uk/biostudies) in MATLAB and Python programming languages under accession number S-BSST641). During this TIF extraction process, each channel image was also max/min rescaled to normalise illumination. Images were also cropped and zero-padded to a standard 64×64 pixel-square size for input into the DeepFlow network.

### Deep learning image classification

Automated scoring was achieved using a nine-class, feed-forward, image classification deep neural network built using our previously described “DeepFlow” architecture (Eulenberg et al. 2017). This network is optimised for the relatively small input dimensions of imaging flow cytometry data, and in itself utilises dual-path convolution / batch normalisation / nonlinearity subunits interspersed by max pooling from the popular “Inception” architecture (Szegedy et al. 2015). These subunit layers process and aggregate visual information at increasing scale before average pooling, the fully connected layer and softmax classification (full network architecture shown, **Supp. Figure 1)**. Images were passed to the network with an input size of 64×64×2 (x, y, channels), with augmentation by random x/y reflection, rotation, translation, 90%-110% image scaling and zero-center batch normalisation. Training lasted for 30 epochs using a batch size of 88 with optimisation under ADAM using cross-entropy loss. The initial learn rate was 5×10^−3^, dropping every five epochs by 0.9, with L2 regularisation 1×10^−4^ and epsilon 1×10^−8^. Images were shuffled every epoch. The final pre-trained network alongside test images and all code detailing training hyper-parameters and final layer weightings are available for download in MATLAB (using the Deep Learning Toolbox) or Python (using TensorFlow / keras) languages at the BioStudies database (http://www.ebi.ac.uk/biostudies) under accession number S-BSST641.

### Ground truth curation by human scoring

For the Cardiff / Cambridge analyses, cell image data across compounds (carbendazim and MMS) and doses (0 – 5 µg/mL) were merged to create diverse ground truth training sets that contained the wide representation of different cell phenotypes essential for effective network training. Ground truth classifications for each image were assigned by biologists with extensive experience manually scoring the *in vitro* micronucleus assay, with phenotypes assigned through consideration of both the nuclear fluorescence and the brightfield image (*i*.*e*., ensuring nuclear events belonged to one cell *etc*.). As per micronucleus assay test guidance, the aim was to only score cells positive for micronucleus events where the micronuclei were fluorescently-labelled, were circular/oval in shape, were within the size range of 1/3 – 1/16^th^ that of the parent nuclei, and that were clearly inside the cell boundary of the parent cell (Fenech 2000; OECD 2016). At the GSK laboratory, TK6 cells were exposed to just the carbendazim compound (0 / 0.8 / 1.2 / 1.6 µg/mL doses) with the experiment conducted in triplicate. For the initial network cross validation with the GSK data, five thousand human-scored cell images were used with these events equally accumulated from across all carbendazim exposures. For the dose-response analysis, cell populations of two thousand events were scored per dose in triplicate by either human-scoring or by the neural network.

### Statistical significance of micronucleus responses relative to control

Assessment of micronucleus response significance was conducted according to the framework described in Johnson et al., (Johnson et al. 2014). Response data was log10 transformed and assessed for normality and variance homogeneity by Shapiro-Wilk and Bartlett tests respectively. Where the transformed data passed these tests (p > 0.05), comparisons of micronucleus responses relative to untreated negative controls employed one sided *post hoc* Dunnett’s test with alpha 0.05. Datasets that failed these tests (p < 0.05) were analysed using the non-parametric *post hoc* Dunn’s test.

### Benchmark dose analysis

To compare the dose-response relationships obtained from human expert scoring relative to those obtained from automatic scoring using the trained neural network, nonlinear regression analysis using the Benchmark Dose (BMD) framework was used. Using the freely available PROAST software, dose-response data were analysed using both the exponential and the Hill model family recommended for the assessment of continuous toxicity data by the European Food Safety Authority (EFSA) (Hardy et al. 2017). In each analysis, combined datasets (*i*.*e*., across scoring methods) were analysed together with ‘scoring method’ specified as a potential covariate (Wills et al. 2016). More complex models with additional parameters were accepted if the fit significantly (p < 0.05; log-likelihood) improved. Here, as in previous work, we found that the log-steepness (*parameter d*) and maximum response (*parameter c*) could reasonably be held equal across dose-response curves, whereas the parameters for background response (*parameter a*), potency (*parameter b*), and within-group variance (*var*) were found to be covariate-dependent (Slob and Setzer 2014). The BMD output describes the ‘equipotent dose’ of the modelled dose-response relationships in addition to the bounding, two-sided 90% confidence interval for each level of the covariate. The benchmark response (BMR) size (also termed the critical effect size) used was 50%, which represents a 50% increase in response relative to the background established in the vehicle (zero-dose) control.

## RESULTS

Here, we investigate the ability of deep learning to provide generalised automation of CBMN assay scoring using imaging flow cytometry data acquired according to local protocols across three different laboratories (Cardiff, Cambridge and GSK). **Fig. 1a** demonstrates our workflow. At the end of the assay, cells were fixed and permeabilised before fluorescent nuclear staining. The choice of nuclear stain varied across the different laboratories according to compatibility with the laser configuration of the local imaging cytometer. At Cambridge, cells were labelled with the blue-fluorescent dye Hoechst 33342 which was stimulated by a 405 nm laser with image capture using a ImageStream^X^ cytometer. At Cardiff and GSK, ImageStream^X^ MKII cytometers were used in conjunction with the red-emitting DRAQ5 nuclear stain and excitation by either a 488 nm or 642 nm laser (respectively). Full details of image acquisition settings at each laboratory are shown in **Supp. Table 1**. Image acquisition speeds depended on cell concentrations, in addition to the time taken to purge the flow stream and load each new sample; approximately ∼ 2000 – 5000 cell-images / minute was typical.

**Fig. 1.**
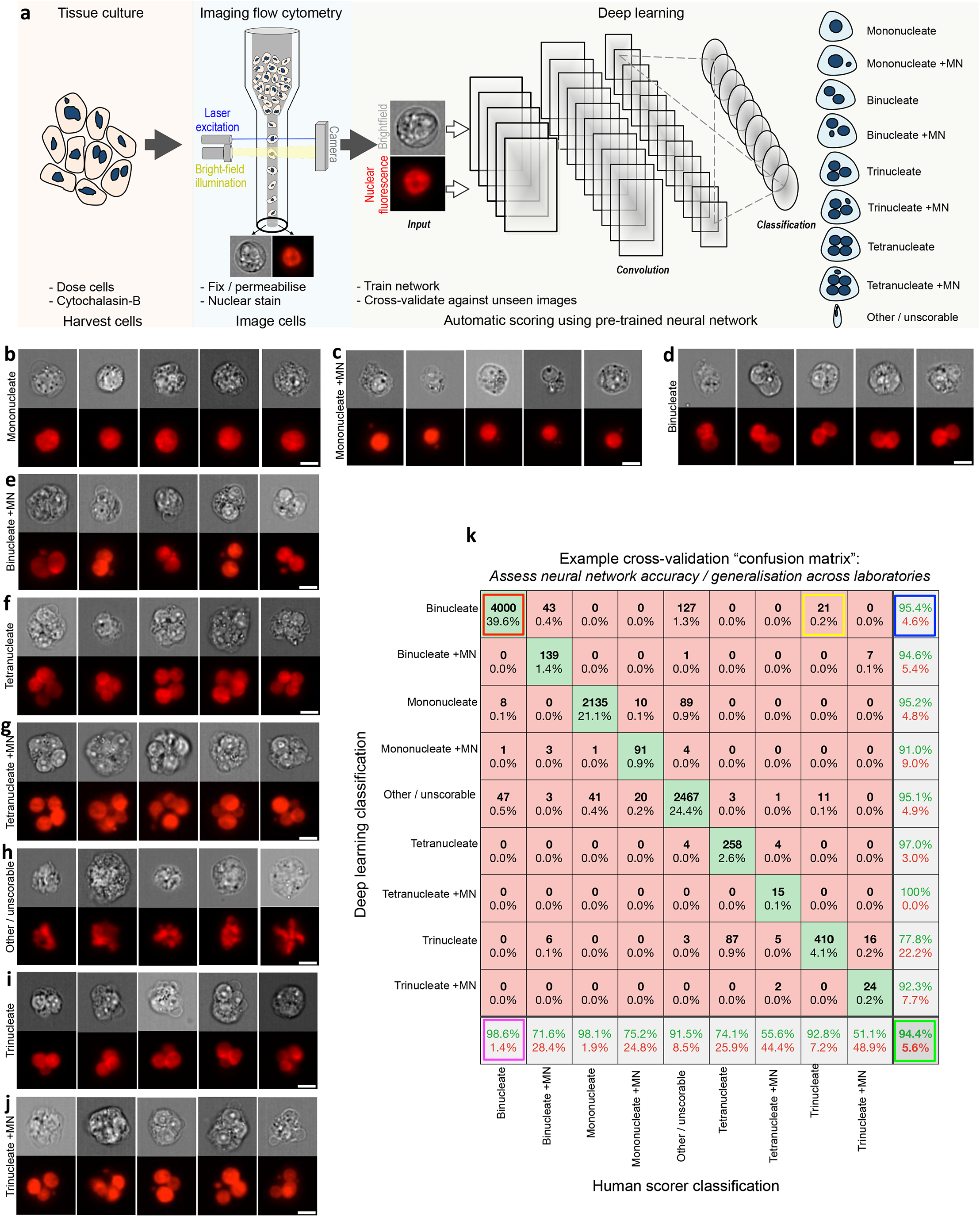
Automating the *in vitro* micronucleus assay using imaging flow cytometry and deep learning image classification. **a** Workflow: harvested cells were fixed and permeabilised before counterstaining the nuclei with a fluorescent DNA stain. Transmitted light brightfield (grey) and nuclear fluorescence (red) images were then automatically captured by high-throughput imaging flow cytometry. After initial training using a human-annotated image set, single cell images from the cytometer can be automatically classified using the neural network image classification algorithm. **b-j** Example image classifications according to a nine-class network developed to score the cytokinesis-block *in vitro* micronucleus assay in human lymphoblastoid TK6 cells. **k** An example cross-validation ‘confusion matrix’ obtained during preliminary network optimisations and presented here to demonstrate confusion matrix interpretation. The matrix represents an image set scored by humans that is ‘unseen’ during network training. The horizontal direction represents the human scorer classifications, whilst the vertical direction shows the automated output classifications from the network. The green diagonal represents correct, matching classifications: for example (indicated, red box) 4,000 ‘binucleate’ images, representing 39.6% of the total test image set, were classified correctly. Away from this diagonal, misclassifications are shown *e*.*g*., (yellow box) 21 images (0.2%) labelled as ‘trinucleates’ by human scoring were incorrectly classified as ‘binucleates’ by the network. In the bottom-right corner (green box) the overall network accuracy and overall misclassification rate are shown for all nine classes (94.4% and 5.6%, respectively). In the white squares down the right-hand side of the matrix, the network precision *i*.*e*., true positive / (true positive plus false positive) (green percentages) and the false discovery rate *i*.*e*., 100-precision (red percentages) are shown for each classification. The horizontal bottom white row shows the network sensitivity *i*.*e*., true positive / (true positive plus false negatives) (green percentages) and false negative rates (red percentages), respectively. Therefore – by example – 95.4% of the images classified as binucleates by the network were binucleates by human-scoring (blue box) whereas the trained model can be expected to correctly assign the binucleate class 98.6% of the time (magenta box). *Scale bars equal 5 microns*

After image collection, a template file created in the cytometer manufacturer’s IDEAS software was used to automatically batch-save populations of single cells that additionally met acceptable focus criteria (see Methods). These cell populations served as the input into the deep learning scoring pipeline. This workflow is provided for download in both MATLAB and Python programming languages at the Biostudies database (accession no. S-BSST641). In brief – the download demonstrates initial image pre-processing to normalise image illumination across cytometers in addition to how to build and train the DeepFlow neural network using a human-scored training image set. After successful training, the saved network can subsequently be used to automate the scoring of new images. For example, **Fig. 1b-j** shows typical events classified by a pretrained, nine-class network with cell classes for mononucleates, binucleates, trinucleates and quadranucleates with or without micronucleus events in addition to a final class for ‘other or unscorable’ phenotypes.

As introduced above, an essential component of network testing involves cross validation with human-scored test images unseen during the training phase. We display this evaluation as a confusion matrix, which compares network outputs to the human scores for every image in the test set (explained, **Fig. 1k**). In the subsequently presented results, we use this strategy to rigorously test the ability of a range of trained networks to enable automated CBMN assay scoring in both intra- and inter-laboratory contexts. In each instance, human-scored image sets were built from cell events pooled across the available compounds and exposures. This strategy was chosen to maximise the diversity of cellular phenotypes present, as well as to ensure that the rarer, micronucleated phenotypes that predominately manifested at higher exposures were well represented.

First, we tested the ability of a network trained on one laboratory’s data to work well for unseen data from that same laboratory (*i*.*e*., ‘single-laboratory testing’) using imaging flow cytometry data collected at either Cardiff or Cambridge (**Fig. 2**). In this single laboratory context, images were randomly assigned to training (60%) and unseen testing (40%) groups. In both instances, the overall accuracies within this single-laboratory context were very high (91.3% and 90.5% for Cardiff and Cambridge, respectively). However, the compiled test sets were quite imbalanced in terms of the numbers of images per class, with network performance with some of the sparser classifications less well represented by the metric of overall accuracy.

**Fig. 2.**
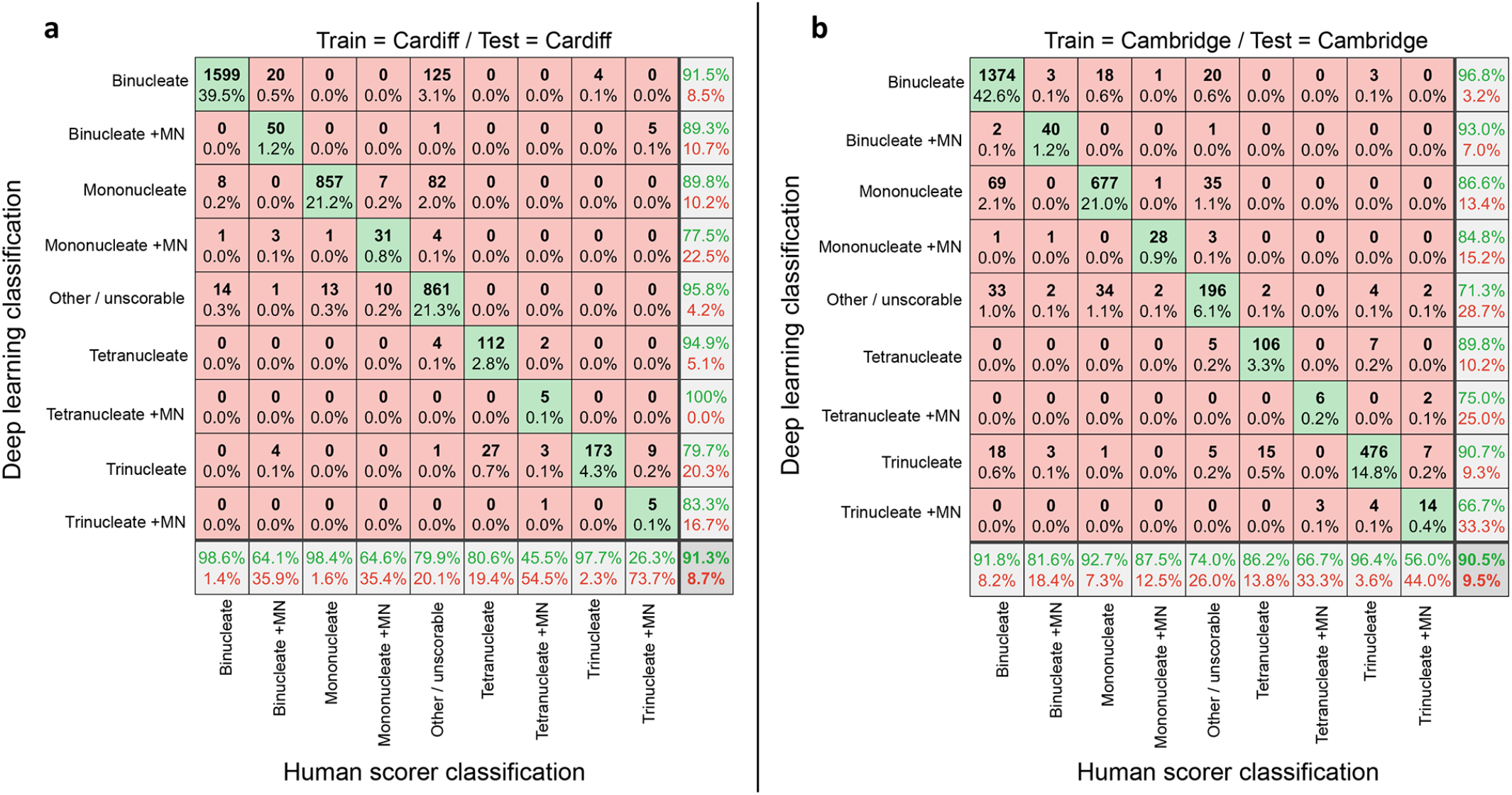
Assessing automated scoring accuracies using intra-laboratory train and test data. **a/b** Confusion matrices comparing human scoring versus deep learning image classifications for test image sets of approximately four thousand unseen images. In each instance, the results reflect the outputs from nine-class networks trained and tested exclusively on image-data from one imaging cytometer at either the **a** Cardiff or **b** Cambridge laboratories

For Cardiff (**Fig. 2a**), whereas accuracy in classification of the common parent nuclei classes (*i*.*e*., mononucleates, binucleates, trinucleates) was generally very good (> 97 %), 20 out of a total of 78 events (∼ 25%) human-scored as ‘binucleate + MN’ were misclassified as ‘binucleates’ by the network. Similarly, around 35% of the human-scored ‘mononucleate + MN’ events were outputted into the ‘mononucleate’ or ‘other/unscorable’ classes, with a further ∼ 20% of ‘tetranucleated’ test images misclassified as ‘trinucleates’. Despite scoring ∼10,000 total events from the Cardiff cytometer, the very rarest cell phenotypes represented by the ‘tetranucleate with MN’ and ‘trinucleate with MN’ classes presented at very low frequency (∼ 0.27 % and 0.47 %, respectively). This led to sparsity in the training set which appeared associated with the network missing micronucleus events, as the ‘trinucleate + MN’ images were often misclassified into the ‘trinucleate’ or ‘tetranucleate’ classes. In a similar manner, ‘tetranucleate + MN’ images were often misclassified into the ‘trinucleate’ or ‘binucleate + MN’ categories.

Similar results were observed within the Cambridge laboratory (**Fig. 2b**). Whereas accuracies with the ‘mononucleate plus MN’ and ‘binucleate plus MN’ classes showed slight improvement when compared against Cardiff, accuracies with the sparser, micronucleated tri- and tetranucleated cells again suffered (∼ 44 and ∼ 33% error rates, respectively).

We next considered the ability of the networks trained on single-laboratory data to generalise to the task of scoring the image data collected from the opposite Centre (**Fig. 3**). This was expected to be a difficult task given that the networks had been trained initially with fairly small numbers of images and because the two laboratories had utilised different cytometer models (IS^X^ vs. IS^X^ Mk II) and nuclear stains (Hoechst at Cambridge or DRAQ5 at Cardiff). This presented the likelihood of overfitting during training – yielding networks highly adapted to the task of scoring data from that particular laboratory.

**Fig. 3.**
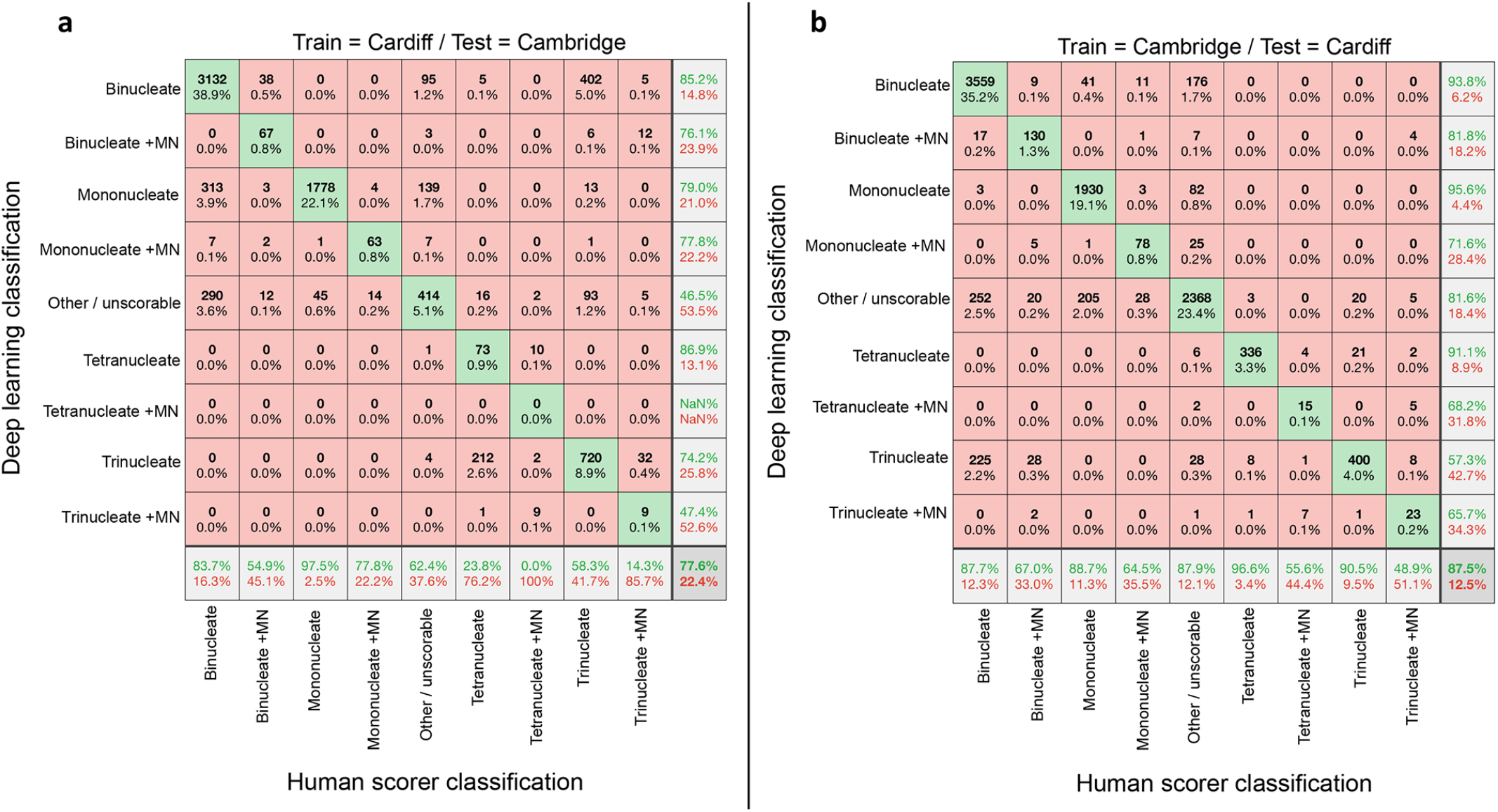
Assessment of automated network scoring accuracies using inter-laboratory test data. **a/b** Confusion matrices comparing human scoring versus deep learning image classifications for test image sets of approximately ten thousand unseen images. In each instance, the results reflect the outputs from nine-class networks trained exclusively on image data from one laboratory’s imaging cytometer before cross-validation testing against image data collected at a different laboratory. **a** Network accuracies after training using Cardiff data before testing on unseen Cambridge data. **b** Network accuracies after training on Cambridge data then testing on unseen Cardiff data

Despite these factors, at first-glance the overall accuracies appeared quite encouraging at 77.6% for the Cardiff-trained network classifying the Cambridge images (**Fig. 3a**) and 87.5% for the Cambridge network classifying Cardiff images (**Fig. 3b**). Comparing across the individual classes, it was apparent that the Cambridge-trained model generalised slightly better to the task of scoring the Cardiff data than was observed *vice-versa*. Closer examination however showed that the metric of overall accuracy was weighted by the prevalence of the easily identified ‘mononucleate’ and ‘binucleate’ phenotypes, which masked assessment of the ability of the networks to identify the micronucleated classes representing DNA-damage events (**Fig. 3a/b**). In this regard, in almost all instances, the accuracy of micronucleated event detection suffered considerably compared to the results achieved with laboratory-matched test data (**Fig. 2**).

With these single-laboratory results established, the images from Cambridge and Cardiff were combined together. This increased the diversity of training exemplifications considerably given the use of two different nuclear stains, two compounds, different imaging cytometers and no ‘hold out’ requirement for cross validation testing. Training a new DeepFlow neural network on this combined training set (∼ 19,000 images) took approximately one hour using modest hardware (single RTX 2080 GPU). The resulting network was then cross validated using a test set where both the bioassay and imaging cytometry were conducted at an entirely new, third laboratory (GSK). Scoring ∼ 5,000 test-images took around six seconds on the RTX 2080 hardware or ∼ 82 seconds on a single CPU. This time, the network showed much better ability to generalise to the task of successfully scoring the images from the new laboratory (**Fig. 4a**). Across the four core classes central to utilisation of CBMN assay (*i*.*e*., ‘mononucleate’, ‘mononucleate plus MN’, ‘binucleate’ and ‘binucleate plus MN’), and with no user input or configuration required, the network achieved 98%, 82%, 94%, and 85% accuracies, respectively.

**Fig. 4.**
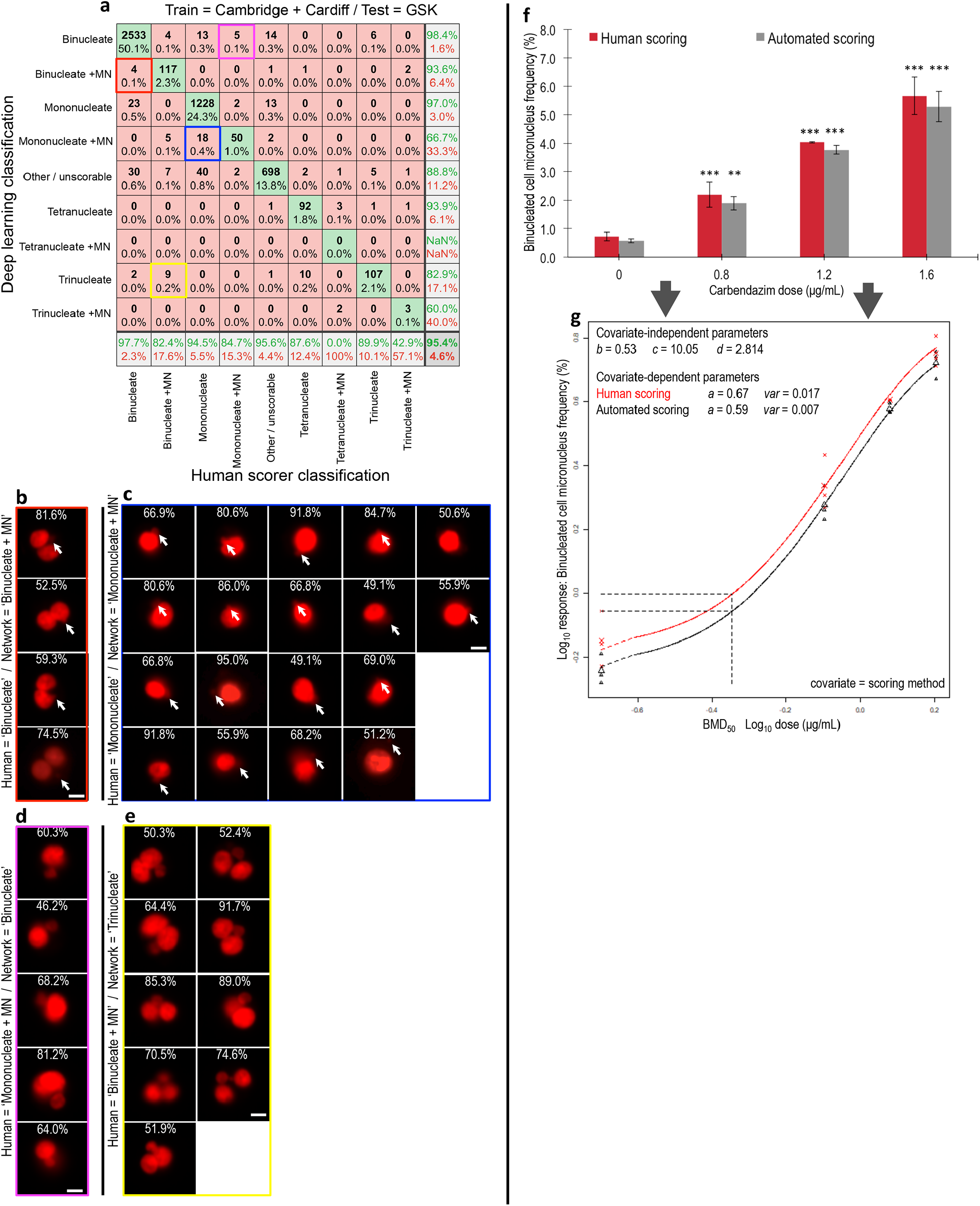
Network accuracy and dose-response assessment using unseen test data from a new laboratory. **a** Confusion matrix showing human versus deep learning image classifications for a test image set of approximately five thousand unseen images. Here, the neural network was trained using image data from both the Cambridge and Cardiff laboratories before testing on new, unseen imaging cytometry data acquired at a third laboratory (GSK). **b** Cell events human scored as ‘binucleates’ but classified as ‘binucleate plus MN’ by the neural network (*i*.*e*., red square in **A**). **c** Cell events human scored as ‘mononucleates’ but classified as ‘mononucleate with MN’ by the neural network (*i*.*e*., blue square in **a**). **b/c**, Close examination of the purportedly misclassified cells shows that many display indistinct events that might be micronucleus or nuclear buds missed by the human scorer (indicated, white arrows). **d** Cell events human scored as ‘mononucleate with MN’ but classified as ‘binucleate’ by the neural network (*i*.*e*., magenta square in **a**). **e** Events human scored as ‘binucleate with MN’ classified as ‘trinucleate’ by the neural network (*i*.*e*., yellow square in **a**). **d/e** In both instances, some of the human-scored micronucleus events encroach upon the 1/3 parent nuclei upper-size limitation typically imposed on micronucleus classifications. **b-e** For each event, the white percentages represent neural network confidence in the outputted classification. **f** Binucleated-cell micronucleus frequencies for a three dose plus control dose-response experiment performed in triplicate for carbendazim exposure to TK6 cells. Scores were established from image sets of 2,000 events per replicate by human scoring or by the cross-validated network established in (**a**). (*) (**) (***) indicate statistical significance relative to control at p < 0.05, p < 0.01 and p < 0.001 respectively. **g** Covariate benchmark (BMD) dose modelling using dose-response data from either the human (black) or automated neural network (red) scores established in (**f**). The horizontal and vertical dashed lines represent interpolation to determine the equipotent, benchmark dose for a benchmark response size of 50%. Regardless of human or automated scoring, the model predicts the same benchmark dose. *Scale bars equal 5 microns*

We then examined failure cases, starting with 22 instances where the network detected micronucleus events in cells scored by humans as just mono-or binucleated (**Fig. 4a**). Surprisingly, many did, in fact, appear to have faint or partially occluded potential micronucleus or nuclear bud events that would have been extremely difficult for the human scorer to detect (**Fig. 4b/c**). Similarly, visualisation of cell events scored by humans as either ‘mononucleate with MN’ or ‘binucleate with MN’, but outputted by the network as ‘binucleate’ or ‘trinucleate’ showed that these images often contained very large micronucleus events (**Fig. 4d/e**). Indeed, some of these likely exceeded the upper size limitation typically imposed on micronucleus classifications (*i*.*e*., ≤ 1/3 diameter of the parent-nuclei) suggesting additional validity to the network’s outputs.

Progressing towards the less frequent cell phenotypes, the accuracies achieved with the ‘trinucleate’ and ‘tetranucleate’ cell classes were also good at 90% and 88% respectively. However, detection of these cell types with micronucleus events was either quite poor or failed entirely. Again, this outcome was likely related to extreme sparsity in occurrence (< 0.25 % frequency in the training data). In an attempt to improve accuracies with these classes, we tried both class weighting the classification layer and combining tri- and tetranucleated events with and without micronucleus events into a single, ‘polynucleated’ class (**Supp. Figure 2**). Whereas both strategies somewhat improved the classification accuracies with these rare events, they were also found to compromise the accuracies achieved with one or more of the four core phenotypes more central to successful CBMN assay scoring.

Given that the frequency of binucleated cells with or without micronucleus events represents the core readout for successful DNA damage assessment by the CBMN assay, after validating the network we proceeded to assess the binucleated-cell micronucleus frequency for a three dose plus control experiment conducted in triplicate with carbendazim at the GSK laboratory. For each dose and replicate, 2000 cell images were scored both manually and automatically. Visually, the resultant dose-response relationships appeared similar across the human and neural network scoring approaches, with the human scores consistently fractionally higher for each dose-group (**Fig. 4f**). To better understand the consequences of this using a recognised, quantitative framework for genotoxic potency estimation, the dose-response relationships were fitted using both the exponential and the Hill model families recommended for the assessment of continuous toxicity data using Benchmark Dose (BMD) analysis (Hardy et al. 2017). With scoring method specified as a potential covariate, model fitting with the PROAST package resulted in covariate-dependent parameterisation for the background response (parameter *a*) and for within-group variation (*var*). For both model families, this parameterisation subsequently allowed rejection of scoring method as covariate, yielding the *same* estimation for the equipotent, benchmark dose from both manual and automated methods (**Fig. 4g**). Model fits to the data are presented in **Supp. Figure 3**.

## DISCUSSION

The CBMN assay represents a globally significant method for the identification and quantification of chromosomal damage (Fenech 2000; Fenech 2020; OECD 2016). Its utility reaches beyond regulatory compound screening to encompass inter-individual monitoring of wide-ranging lifestyle, occupational and environmental factors (Fenech 2020; Kirsch-Volders et al. 2011; Wang et al. 2019). Despite this, continued reliance upon time-consuming and user-subjective manual scoring represents a bottleneck to broadening practical utilisation (Seager et al. 2014; Verma et al. 2018; Verma et al. 2017). In this pilot study, we show that rapid image acquisition by imaging flow cytometry in conjunction with deep learning image classification represents a capable platform for automated, inter-laboratory operation. We share our strategy via openly accessible frameworks.

As an image acquisition method, imaging flow cytometry is now well established as a means for high-throughput CBMN data capture with concomitant image archiving potential (Rodrigues et al. 2014a; Rodrigues et al. 2016a; Rodrigues et al. 2018). Moreover, this is achieved with simple sample preparation involving a single nuclear stain and brightfield to provide the context that events lie inside parent cells (Rodrigues et al. 2018). Comparison studies have shown that the captured images contain dose-response information that aligns to results obtained from ‘gold standard’ manual microscopy scoring (Verma et al. 2018). Whereas conventional flow cytometry offers faster throughput, it lacks this image-based validation whilst additionally requiring cell lysis. This prevents utilisation of the cytokinesis-block version of the assay with its associated advantages such as robust utilisation of primary human cell lines, knowledge that cells have divided during the test period and quantitation of mononucleated, binucleated and different classes of multinucleated cells. This information is useful in the avoidance of misleading negative results and additionally enables calculation of division and replication indexes that contribute to assessments of mitogen response and cytostatic impact (Rodrigues et al. 2018).

Beyond image collection, automated scoring of imaging flow cytometry data – as with other automated microscopy strategies – has thus far largely relied upon traditional, threshold-based image classification techniques. These require image analysis expertise to implement, alongside user- configuration and tuning to maintain performance (Rodrigues et al. 2018; Seager et al. 2014; Verma et al. 2017). Unfortunately, much as with traditional manual scoring, this is time-consuming and subjective.

In contrast, once successfully trained, the results achieved here suggest that deep learning image classification has the potential to eliminate these expertise and user-input requirements, dramatically reducing the time to results. This comes from encompassing image diversity during network training and harnessing it to improve the consistency and robustness of subsequent classifications. To this end, here we show that utilisation of diverse training data curated across two laboratories utilising different nuclear stains, multiple compounds and two different cytometer models yielded a capable neural network for scoring automation. Without user configuration, the network was able to classify data collected from an entirely new laboratory with > 82% accuracy for each of the four cell phenotypes central to CBMN performance (*i*.*e*., mononucleate and binucleate cells with or without micronucleus events) in addition to successfully classifying tri- and tetranucleated cells (> 88% accuracy) and unscorable events (96% accuracy). Importantly, these seven classes encompassed virtually all of the cell images encountered (>99%). Success at micronucleus detection in both mononucleate and binucleate cell classes further suggests that this single network could be used to automate scoring of both mononuclear and cytokinesis-block versions of the assay.

Despite this success with the assay classes central to CBMN scoring, the scarce, tri- and tetranucleated phenotypes with micronucleus events proved more challenging. Commonly employed methods such as class weighting or class combination offered little in the way of accuracy improvements, and often compromised accuracy with the other classes. These findings suggest that significant increases in the representation of these sparse events during training will likely be required to improve success. In this context, imaging flow cytometry is well suited to examine whether an improved image bank leads to enhanced accuracy in scoring given the high rates of image capture achievable. Our results also suggests that class reduction does not necessarily simplify the classification problem and may instead cause ambiguities. In this way, future expansions to the number of classes to encompass all distinctive cellular phenotypes may represent a route to improving overall network performance.

In this regard, we identified additional, potentially-scorable cell phenotypes (**Fig. 5**). In particular, cell death events (*i*.*e*., due to apoptosis and necrosis) were visually apparent, but we were unable to determine apoptotic from necrotic events using just the brightfield and nuclear fluorescence images alone. Cells caught during mitosis also represented distinctive events. At the same time, we were less convinced that more subtle phenotypes relevant to the expanded, CBMN cytome assay such as nuclear buds and bridges could reliably and consistently be detected – given the relatively low resolution of the image data (Fenech 2007). However, it is important to note that previous studies demonstrating capture of these phenotypes by imaging flow cytometry have utilised both the 60X ImageStream objective lens in addition to hypotonic treatments to swell cell volumes prior to imaging (Rodrigues 2019; Rodrigues et al. 2018). Hypotonic treatments were not used here but may improve image capture of these more subtle phenotypes. With regards to network class expansion to encompass these events – or, indeed for simultaneous measurement of other endpoints – the ImageStream platform is capable of multiplexed imaging. Additional channels might therefore be used to simultaneously measure other DNA-damage pathways (*e*.*g*., ϒH2AX for DNA double-strand breaks (Smart et al. 2011)), or to improve the reliability of ground truth image curations through use of additional fluorescent markers to differentiate events such as apoptotic from necrotic cells.

**Fig. 5.**
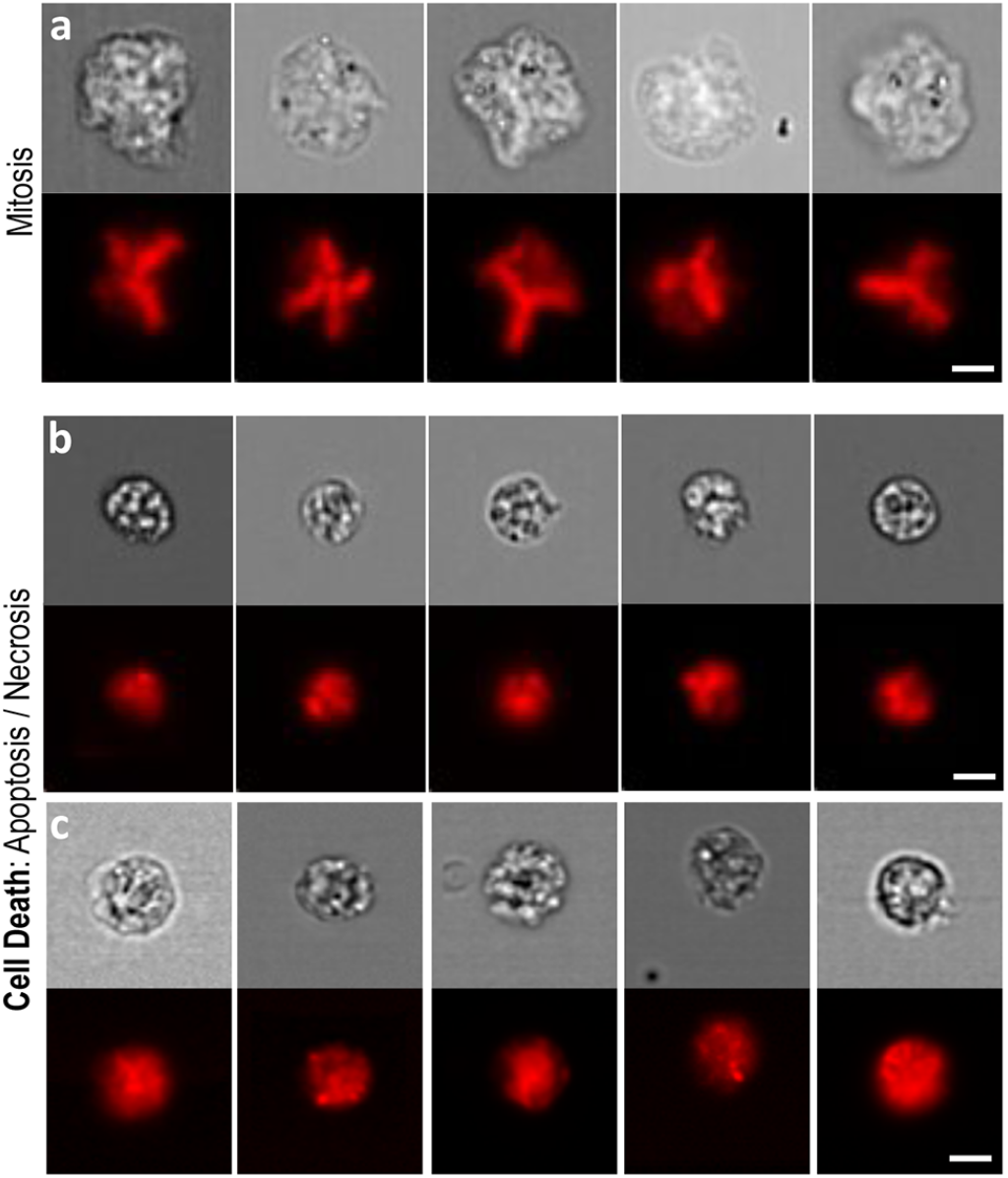
Other scorable cell phenotypes captured by imaging flow cytometry. **a** Cells undergoing mitosis were visually apparent according to metaphase spread-type nuclear fluorescence imagery (red) alongside large, brightfield-delineated cell sizes (grey). **b/c** Cell death events displayed shrunken cell sizes in conjunction with granular brightfield and fluorescence imagery. In the case of cell death, two distinctive cell phenotypes appeared visually separable according to cell size and the number, size and extent of nuclear foci formation (**b** versus **c**). Whether these observations represented distinct apoptotic versus necrotic events was unclear from the nuclear fluorescence and brightfield information alone. *Scale bars equal 5 microns*

Manual scoring of the images for this experiment was more challenging than the exemplar images shown might suggest. Fundamentally, the acquired images are relatively low resolution (*i*.*e*., cells occupy ∼ 64×64 pixels) and further image degradation is always present as a result of the capture of moving objects by time delay integration. The acquired images also represent a central, 2-D projection of a 3-D cell-object. This means that nuclei and micronucleus events may overlap each other, or they may lie outside of the plane of optimal focus (Rodrigues et al. 2018). These factors all served to make ground truth assignments more complicated, even for experienced CBMN scorers. Whereas network accuracy assessments by confusion matrix provided a more representative breakdown of outputs when compared to simplistic overall accuracy measures, it is a relatively stringent success measure because any ambiguity in human score assignment is not captured. A potential advantage of automated network classification approach is therefore likely greater consistency – even in error – than arises from manual scoring.

Regarding image focussing, the ImageStream platform offers ‘extended depth of field’ (EDF) technology, whereby image deconvolution is used to improve the utility of out of focus events through projection onto a single plane (Ortyn et al.). Whereas previous studies have shown this technique can improve accuracy in ‘spot counting’ applications, the strategy has been reported less helpful for the provision of improved CBMN data (Parris et al. ; Rodrigues 2018; Rodrigues et al. 2014a). This was attributed to a slight degradation in overall image resolution, compromising differentiation of micronucleus events from parent nuclei (Rodrigues 2018). On a similar theme, the ImageStream platform is also configurable with 20X, 40X or 60X objective lenses. Here, image collection was via the ‘standard’, 40X objective across all laboratories. This approach was chosen as previous work has shown that whilst greater resolution is achievable with the 60X objective, focus depth also decreases, reinforcing the out of plane difficulties described above (Rodrigues et al. 2018).

Whilst considering the nature and utility of imaging flow cytometry data, a relevant comparison is to that provided by other automated imaging methods such as slide scanning platforms. In addition to the potential for higher resolution imaging, here an overlooked advantage comes from the ability to use slide-based preparations created by cytocentrifugation. This technique causes the flattening and spreading of cellular content, presenting nuclear objects on a more two dimensional plane (Fitzgerald and Hosking 1982; Shanholtzer et al. 1982). From a practical perspective however, this also necessitates the consistent preparation of high-quality slides with optimal cell densities (Rodrigues et al. 2018). Meanwhile, a major advantage of the imaging flow cytometry approach is that single cell image data is inherently acquired by the fluidics-based processing of individualised cells.

## CONCLUSIONS

As a platform for the CBMN assay, imaging flow cytometry combines the high throughput and multiplexing potential of flow cytometry with the image-based validation and archiving attributes of automated microscopy. Here we demonstrate accurate, automated assay scoring using a neural network for data collected in a laboratory wholly separate to that in which the algorithm was trained. This proves that without any human configuration, the machine is able to correctly anticipate the decisions of the expert human on unseen images in a new setting. For the first time, this suggests the possibility for generalised scoring automation through dissemination of a pretrained network for the ImageStream platform established from ground truth agreed by a single, expert group. Such an approach would provide the ultimate in terms of standardisation and result reliability, but more importantly could enable adoption of the assay beyond current practitioners as local expertise in scoring and/or image analysis would no longer be required. For these reasons, we believe that full development of this automated, accessible, inter-laboratory approach would represent a truly twenty-first century method with significant potential to transform CBMN utility across industry, research and clinical domains.

## Acknowledgements

The authors thank Dr. R. Wilkins and Dr. L. Beaton-Green at Health Canada for sharing their expertise. J.W.W. is grateful to Girton College and the University of Cambridge Herchel-Smith Fund for supporting him with fellowships.

---

**Supp. Table 1–.** Imaging flow cytometry data acquisition information.

| **Centre** | **Excitation laser (nm)** | **Intensity (mW)** | **Brightfield channel** | **Nuclear fluorescence channel** | **Nuclear stain** | **Objective lens** | **Cytometer Model** |
| --- | --- | --- | --- | --- | --- | --- | --- |
| Cambridge | 405 | 50 | Ch04 | Ch01 | Hoechst 33342 | 40X | Amnis ImageStream<sup>x</sup> |
| Cardiff | 488 | 100 | Ch01 | Ch11 | DRAQ5 | 40X | Amnis ImageStream<sup>x</sup> MkII |
| GSK | 642 | 55 | Ch01 | Ch11 | DRAQ5 | 40X | Amnis ImageStream<sup>x</sup> MkII |

Image data were collected using three different imaging flow cytometers located across three laboratories (Cambridge, Cardiff and GSK). At each laboratory, the choice of fluorescent nuclear stain depended upon local protocols and compatibility with the cytometer's laser configuration.

**Supp. Figure 1.**
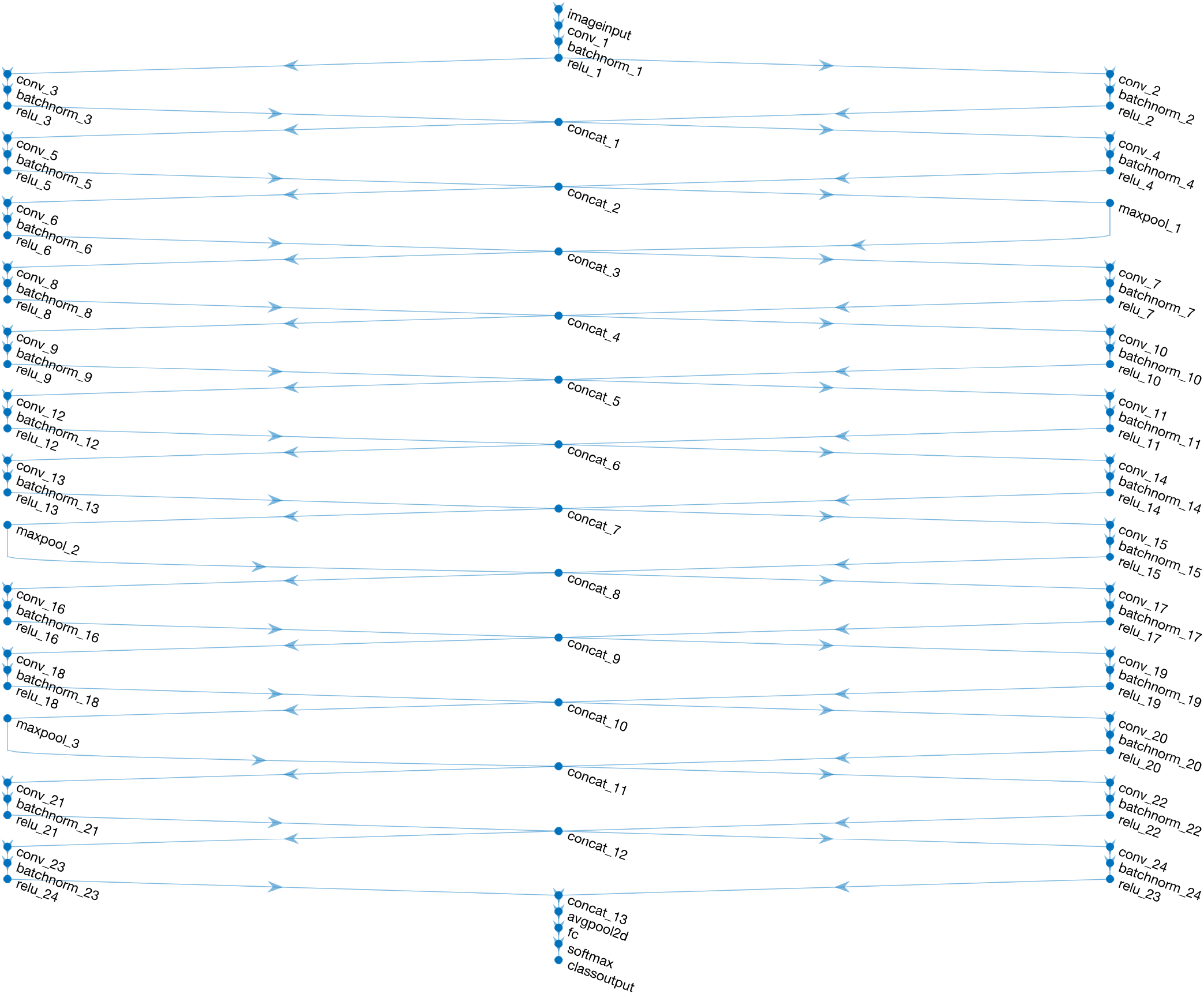
DeepFlow neural network architecture schematic. The DeepFlow network utilises a 64×64×2 input layer (x, y, channels) followed by repeating dual-path subunits from the “Inception” architecture to aggregate visual information over increasing scales. The number of kernels used increases at each layer, yielding 336 features maps with size 8 x 8 before average pooling, the fully connected (fc) layer and softmax classification using cross-entropy loss.

**Supp. Figure 2.**
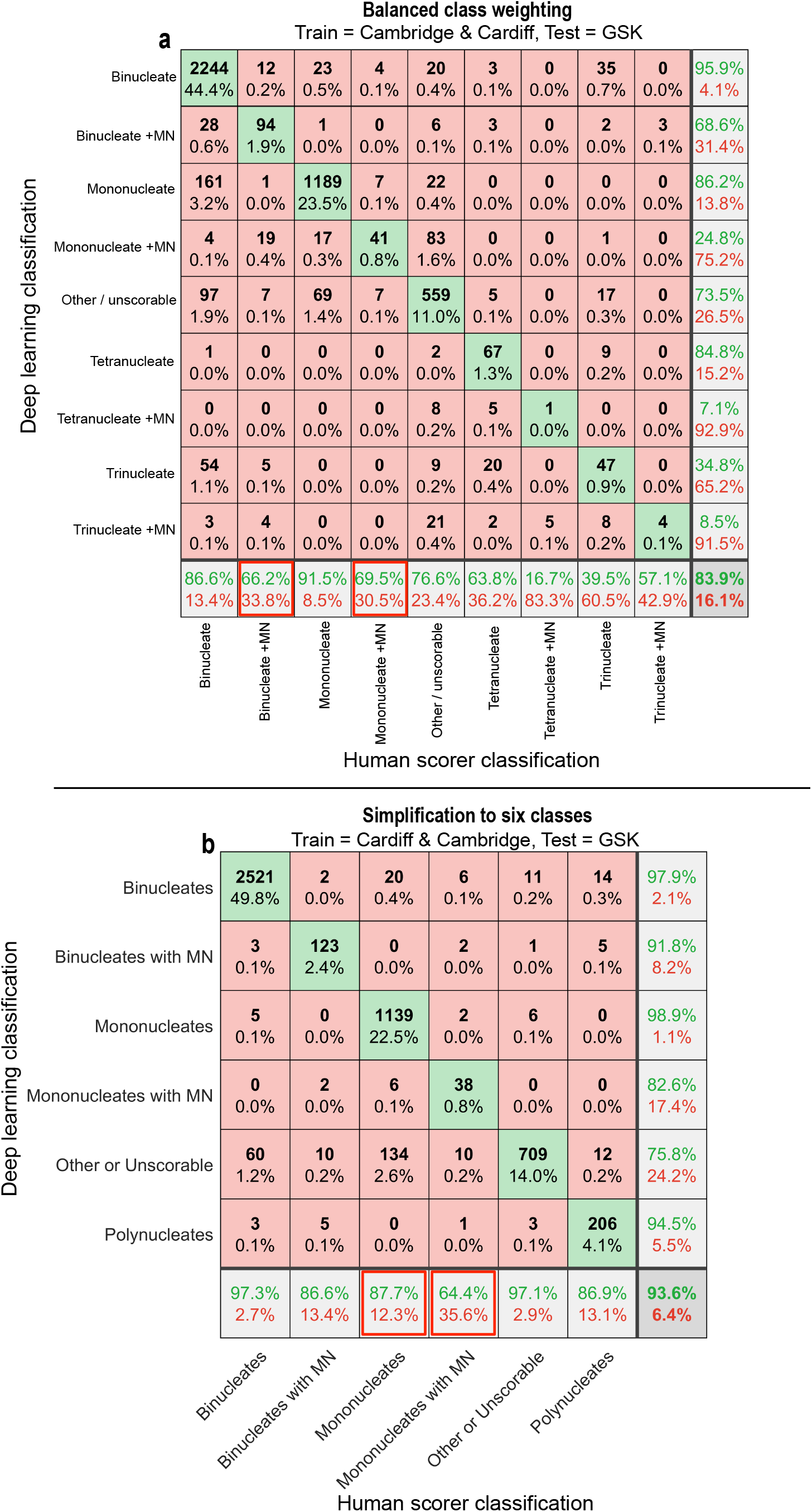
Cross validation testing using class weighting or class simplification strategies. **a/b** Confusion matrices comparing human scoring versus deep learning image classifications for a test set of ∼ 5000 unseen images. In each instance, the results reflect the outputs after training using image data from both the Cambridge and Cardiff laboratories before cross validation on new imaging cytometry data acquired at a third laboratory (GSK). In **a** class weighted cross entropy loss was used at the classification layer in an attempt to improve performance with the sparsely-represented phenotypes (*i*.*e*., tri and tetranucleates with or without micronucleus (MN) events). In **b** these sparse, multinucleated categories were combined together into a single ‘polynucleated’ class. Whilst some improvements were realised using these strategies, they both reduced achieved accuracies (indicated, red squares) with one or more of the four, core phenotypes central to successful CBMN scoring (*i*.*e*., mono or binucleated cells with or without MN events).

**Supp. Figure 3.**
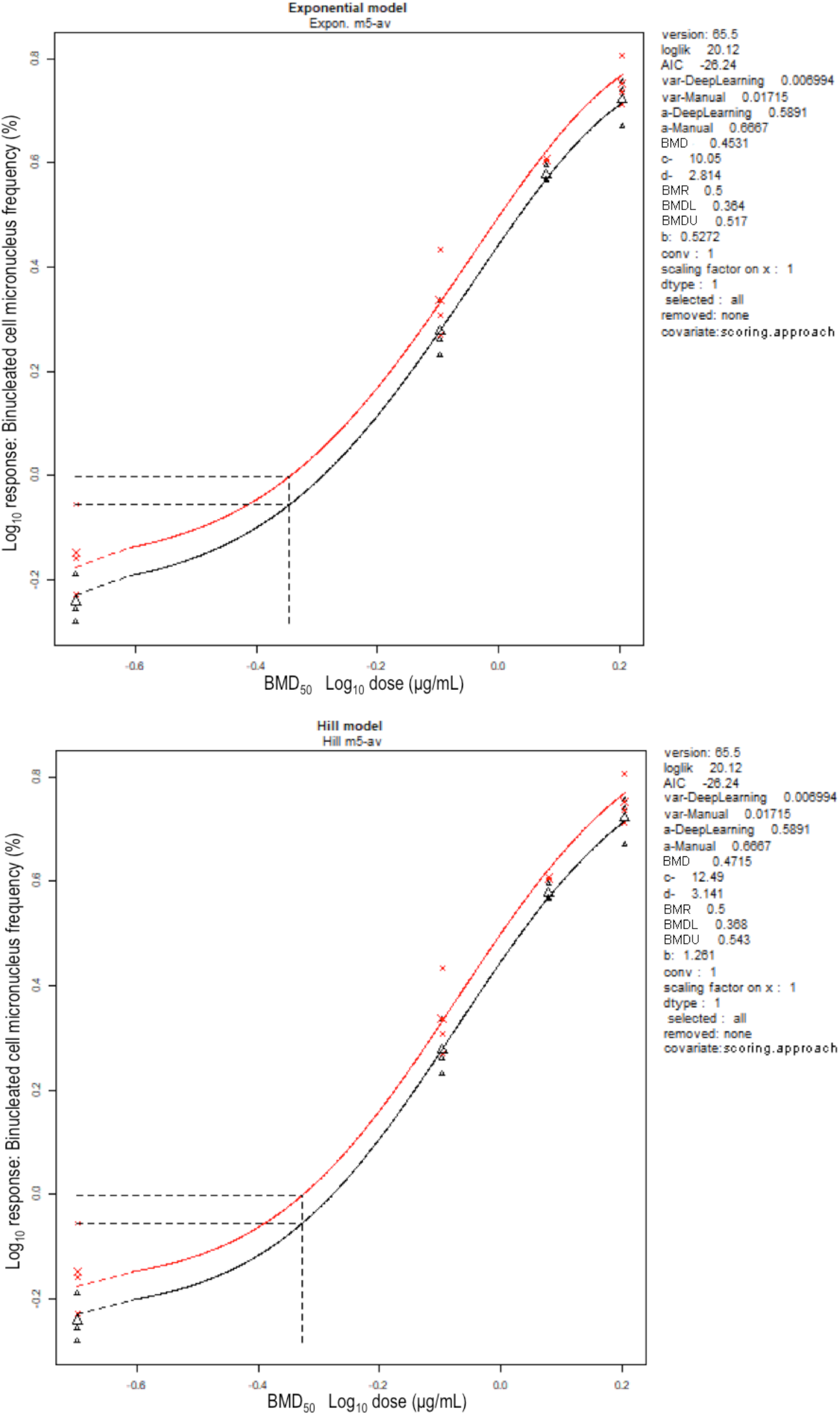
Benchmark dose (BMD) analysis using exponential and Hill model families. The curves represent fits to micronucleus dose-response data obtained either by human (red) or neural network (black) scoring using either the exponential (top) or the Hill (bottom) model families. Both models were fitted with covariate (scoring method) dependent parameters for the background (parameter *a*) and within-group variance (*var*), whilst constant parameters could be used for potency, shape and steepness (parameters *b, c* and *d*). Horizontal and vertical dashed lines represent interpolation at a benchmark response (BMR) size of 50% to determine the BMD50 (respectively).

## Notes

**Conflicts of interest/competing interests** M. A. R., is an employee of Luminex Corporation which manufactures the Amnis ImageStream imaging flow cytometers used in this research study.

https://www.ebi.ac.uk/biostudies/studies/S-BSST641

### Competing Interest Statement

The author M. A. R., is an employee of Luminex Corporation which manufactures the Amnis ImageStream imaging flow cytometers used in this research study.

